# Pumping Up your Predictive Power for Cognitive State Detection with the Proper GAINS

**DOI:** 10.1101/2025.04.01.646685

**Authors:** Victoria Ribeiro Rodrigues, Jeremy R. Prieto, Szilard L. Beres, Chad Stephens, Christopher Myers, Nicholas J. Napoli

## Abstract

Detecting cognitive states and impairments through EEG signals is crucial for applications in aviation and medicine and has broad applications in the field of human-machine interaction. However, existing methods often fail to capture the fine-grained neural dynamics of critical brain processes due to limited temporal resolution and inadequate signal decomposition techniques. To address this, we introduce the Spectral Intensity Stability (SIS) algorithm, a novel technique that analyzes the stability and competition of dominant brain frequency oscillations across granular timescales (4 ms). Unlike traditional spectral methods, SIS captures rapid neural transitions and hierarchical frequency dynamics, enabling more accurate characterization of task-specific cognitive processes. Our study focuses on EEG data from pilots performing multitasking simulations under hypoxic and non-hypoxic conditions, a high-stakes scenario where cognitive performance is crucial. We divided this multitasking scenario into specific cognitive states, such as task precursor, interruption, execution, and recovery. Our algorithm SIS achieved a 29.8% improvement in cognitive state classification compared to conventional methods, demonstrating superior accuracy in distinguishing both task states and hypoxic impairments. This work is novel because it bridges gaps left by traditional methods by revealing the role of hierarchical spectral dynamics in maintaining cognitive performance. Through the Granular Analysis Informing Neural Stability (GAINS) framework, we reveal how neuronal groups self-organize across fine-grained time scales, providing new understanding of task-switching, neural communication, and criticality. The findings highlight the potential for developing real-time cognitive monitoring systems to enhance safety and performance in environments where cognitive impairments can have serious consequences. Future research should extend these insights by incorporating transient behaviors and spatial dynamics to achieve a more comprehensive framework for characterizing cognitive states.

## 1. Introduction

Electroencephalogram (EEG) signals capture information about electrical activity in the human brain, which has been widely used to detect and classify cognitive states and cognitive impairments. However, current state detection methods often overlook the rich information contained within oscillatory brain activity (neural oscillations) due to limitations in existing signal processing tools and techniques. Neural oscillations, categorized as delta (0–3.5 Hz), theta (4–7 Hz), alpha (8–12 Hz), beta (13–40 Hz), and gamma (32–100 Hz) bands, encode information about the synchronization of neural firing, which facilitates communication within and between brain regions. While this communication and its relationship to cognitive states are well understood, the underlying neural mechanisms driving cognitive functionality remain unclear [10, 12].

Capturing and characterizing these underlying mechanisms and processes could enable the development of more advanced cognitive state detection systems, ultimately enhancing human-machine interaction. Therefore, it is imperative to improve signal processing tools and techniques to better analyze and decode the dynamics and neural mechanisms underlying cognition.

### 1.1. Prior Work

To better understand the dynamics underlying cognition and cognitive processes, it is essential to develop methods for extracting information from human brain activity, such as EEG signals. This study explores the underlying dynamics of how multiple neural oscillatory groups organize to fire together, leading to the emergence of new groups or the inhibition of competing neural oscillations. These organizational transitions can be described as neural criticality, a phenomenon where a system undergoes a phase transition [35, 16, 4]. While the detailed mechanisms of neural criticality and brain activity dynamics remain only partially understood, previous research suggests that healthy brain activity operates near critical phase transitions. These boundaries, often referred to as critical points, delineate synchronized and asynchronous neural activity [35, 12, 22, 48].

We hypothesize that these critical points are fundamental to the brain’s information-processing capabilities and that different cognitive tasks may rely on distinct types of organizational processes. These processes facilitate more effective communication between neural oscillator groups, giving rise to unique cognitive functions [35, 11, 12, 41]. *This raises a fundamental question: If different types of organizational processes are conditioned for specific cognitive functions, how can we decode EEG signals to appropriately characterize these processes in a way that reflects their respective critical points?*

#### Human Activity & Criticality Across Timescales

To begin addressing this fundamental research question, we draw from Newell [28], who discusses the various time spans across different time scales that define the range of human activities, from the biological to the social band. This framework provides the basis for a proper *granular analysis* necessary to uncover how neuronal groups organize and interact. Human activities can be characterized across time scales spanning 12 orders of magnitude, from processes occurring within organelles (i.e., the biological band; 10^−4^ s) to activities within and across societies (i.e., the social band; 10^7^ s) [see Table 1].

**Table 1:** Newell’s Time Scale of Human Action.

| Time Scale (secs) | Human Action | Newell’s Band |
| --- | --- | --- |
| $10^7$ (months) | Technology | Social |
| $10^6$ (weeks) | Design | Social |
| $10^5$ (days) | Task | Social |
| $10^4$ (hours) | Task | Rational |
| $10^3$ (10 minutes) | Task | Rational |
| $10^2$ (minutes) | Task | Rational |
| $10^1$ (10 seconds) | Unit Task | Cognitive |
| $10^0$ (second) | Operations | Cognitive |
| $10^{-1}$ (100 ms) | Deliberate Act | Cognitive |
| $10^{-2}$ (10 ms) | Neural Circuit | Biological |
| $10^{-3}$ (1 ms) | Neuron | Biological |
| $10^{-4}$ (μs) | Organelle | Biological |

According to Newell’s framework, cognitive behaviors often emerge from the novel integration of processes occurring at finer levels of analysis. For example, observable actions performed at the operations time scale (approximately 1 second, such as moving a mouse pointer and clicking on an object; see Figure 1) are driven by cognitive processes occurring at the deliberate acts time scale (approximately 100 ms). Within this hierarchical structure, processes and outcomes at shorter time scales serve as the foundation or inputs for more complex cognitive processes that unfold over longer durations (see Figure 1).

**Figure 1:**
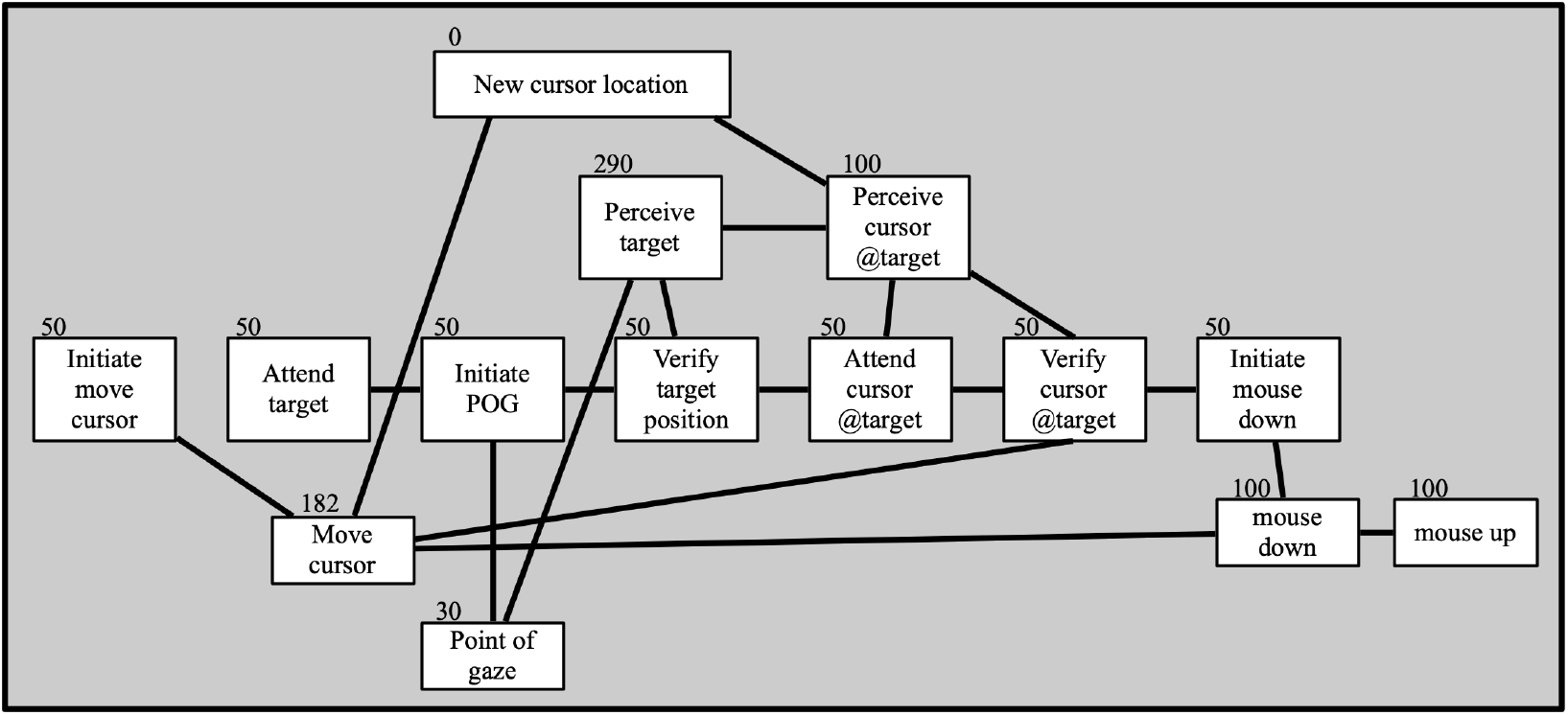
A Goals, Operators, Methods, and Selection Rules (GOMS) model illustrating deliberate acts organized into the operation of moving a mouse pointer to a target and clicking on the target (adapted from [15]). The numbers above each process indicate the process duration in milliseconds, while the lines between the processes represent dependencies. The duration of this operation ranges from 450 ms to 1152 ms.

To enhance cognitive task detection, these finer-grained time scales—such as deliberate acts (10^−1^ seconds) and operations (10^0^ seconds)—should be examined as they contain critical information about the organizational dynamics of neuronal oscillatory groups, specifically the activation of neural criticality associated with underlying cognitive processes. However, based on the principles of criticality and Newell’s defined time bands, current methodologies are insufficient to effectively capture this level of temporal resolution and its dynamics. This creates a significant gap in our ability to decode neuronal firing patterns, cognitive processes, and impairments occurring within these critical granular time scales (100 ms to 1 second).

The concept of criticality, when viewed through the lens of Newell’s time scale, suggests a potential relationship between smaller time scales (i.e., building blocks) that exhibit criticality and contribute to the emergence of larger processes, such as cognition. This self-organization process involves neurons firing synchronously, triggering additional neurons to fire in a coordinated manner, thereby altering the state of the globally recorded neuronal network.

To advance our understanding of neuronal dynamics, it is crucial to develop analytical methods capable of decomposing signals into distinct neuronal oscillatory groups within granular time scales. Such granular analysis would enable a deeper exploration of the interactions and self-organization of these oscillatory groups, providing valuable insights into the mechanisms that underpin the stability of neuronal firing, criticality, and their roles in cognitive processes. Consequently, an enhanced *Granular Analysis Informing Neural Stability (GAINS)* framework could offer the necessary information to effectively discriminate between different cognitive processes.

#### Current Analysis Limitations for State Detection

Traditional EEG signal decomposition methods, such as Fourier analysis, aim to extract a signal’s intensity at specific EEG frequency oscillations (bands) to obtain spectral features, which are then associated with cognitive behaviors, processes, learning, and impairments [47, 3, 50, 19, 36]. The Fast Fourier Transform (FFT) accomplishes this by projecting the time-domain EEG signal into the frequency domain, consisting of its associated frequency and phase components (see the central plot of Figure 2 for the frequency spectrum). However, in analyzing cognition using traditional Fourier approaches, EEG signals can only be resolved using window lengths on the order of seconds (e.g., one second [50, 36], two seconds [17, 47], four seconds [40, 19]). While this allows for blunt characterization of spectral intensity, it neglects information on spectral intensity fluctuations, rendering traditional approaches incapable of capturing events of criticality and the neural communication between oscillatory groups. Figure 2 illustrates this limitation by showing how traditional methods fail to capture events of criticality and changes within oscillatory groups.

**Figure 2:**
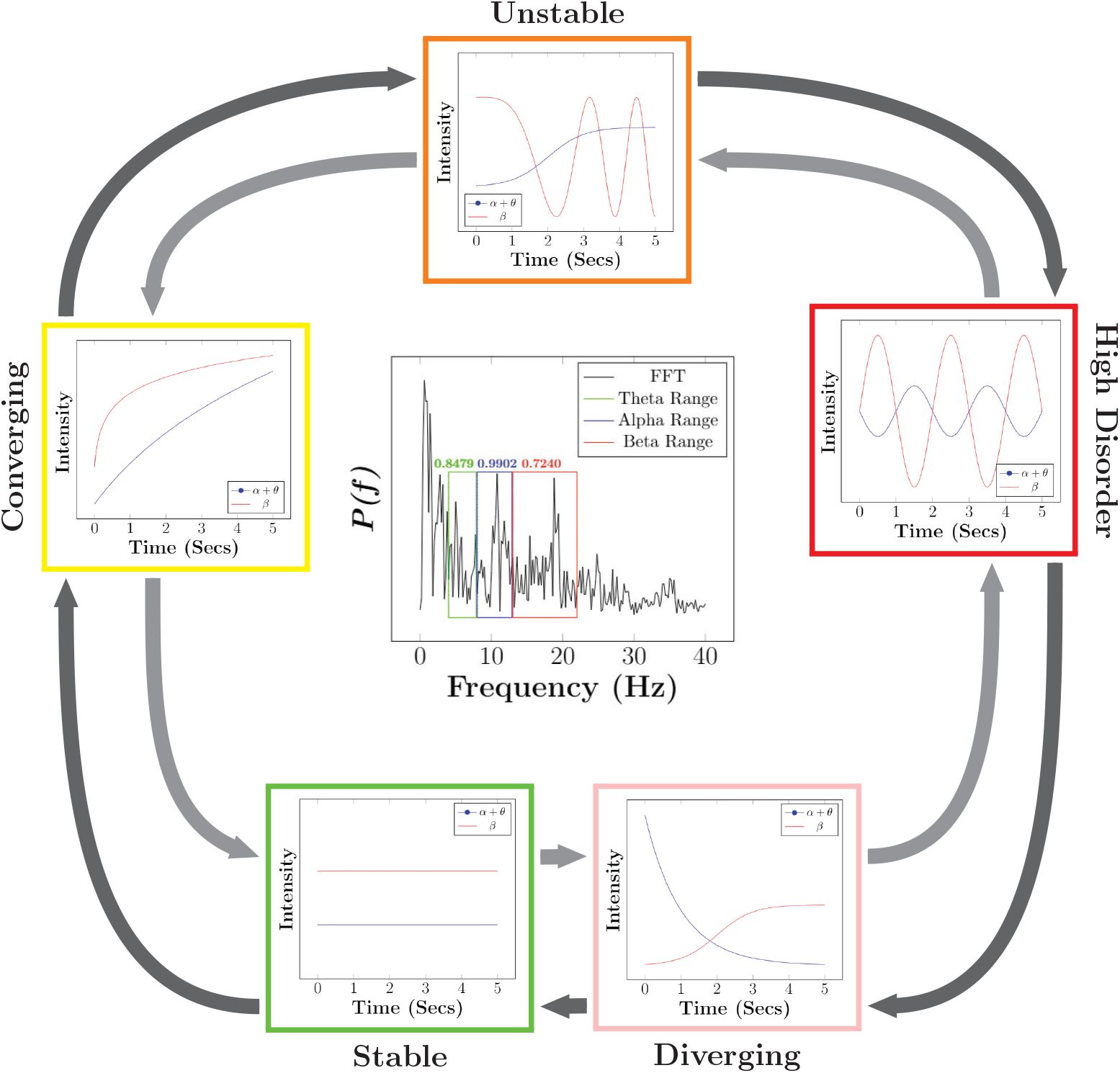
Potential EEG spectral intensity dynamics for characterizing cognitive task transitions are illustrated. These dynamics are broadly categorized into five states: Converging, Unstable, High Complexity, Diverging, and Stable. The arrows depict the possible transitions of these dynamics over time, emphasizing the relevance of Newell’s timescale in decoding cognitive processes. Stability can either increase (gray arrows) or decrease (black arrows). This framework introduces the concept of Spectral Intensity Stability (SIS), where different frequency bands exhibit distinct hierarchical arrangements and stable dynamics, potentially forming a cognitive process or state. Each graph represents a different level of SIS that could occur within a given time window.

The temporal spectral intensity dynamics, as shown in the figures surrounding the central blue plot in Figure 2, can vary significantly over time. However, these dynamics become indistinguishable when the temporal dimension is collapsed, leaving only the frequency content in the central plot. This obscures critical points and dynamics necessary for distinguishing potential cognitive states, such as Unstable, Converging, High Disorder, Stable, and Diverging. Each of these states reflects distinct neuronal firing criticality and neural communication patterns, yet projects the same frequency domain representation. Consequently, traditional frequency spectrum analysis only measures the intensity of frequencies during a time window but fails to capture the dynamics of hierarchical arrangements or the stability of neuronal populations within that period. Periods with vastly different dynamics may produce identical Power Spectral Densities (PSD), providing no information about the hierarchical arrangement or stability of neural oscillatory groups.

This underscores the need to develop novel methods capable of accurately representing cognitive dynamics and their relationship to cognitive processing and states. Current analyses overlook critical temporal scales (such as those defined by Newell’s cognitive bands within unit operations and deliberate acts) and the nature of hierarchical arrangements of neuronal oscillatory groups within a single spatial location. Alternative approaches are required to fully capture the underlying mechanisms and dynamics of criticality associated with cognitive processes.

Existing metrics, such as the Engagement Index (EI) [32], have shown potential in measuring the hierarchical arrangement between neural oscillations and classifying cognitive states. However, these methods lack the granularity needed to characterize dynamics adequately and provide only blunt approximations [32, 33, 5]. Therefore, the development of advanced metrics that quantify the dynamic hierarchical arrangements of neural oscillations within appropriate temporal scales (as defined by Newell’s cognitive bands) could offer significant insights into the criticality underlying a “fingerprint” for specific cognitive processes.

This leads to two fundamental research questions regarding criticality and cognitive processes:

RQ1 Do unique dynamics within these task events that elicit criticality have unique dynamics, potentially establishing a “fingerprint” for cognitive processes?

RQ2 Does the depression of neuronal firing due to hypoxia lead to events of criticality with unique spectral arrangements?

## 1.2. Challenges

To effectively capture critical points and transitions in neural activation within cognitive states, methods capable of extracting both granular spectral and temporal information are essential. Wavelet analysis has emerged as a promising alternative to Fourier methods for identifying instances of criticality [18, 7, 44]. Wavelets allow for the continuous extraction of spectral information over time, offering improved granularity. However, conventional wavelet techniques face resolution limitations that hinder their ability to fully capture the underlying mechanisms of critical points. Wavelets use basis functions scaled in the time domain to create a filter bank in the frequency domain, with unique center frequencies and bandwidths arranged linearly or logarithmically [20, 44, 34]. This scaling leads to overlapping, non-orthogonal wavelets, which reduce the precision of frequency extraction for neuronal oscillations such as delta, theta, alpha, beta, and gamma [24]. Overlapping wavelets result in redundant frequency components and imprecise signal reconstruction, limiting the accurate analysis of neural criticality within oscillatory group communication and hindering a comprehensive understanding of hierarchical frequency oscillation dynamics. Consequently, standard wavelet transformations and time-frequency analysis fall short of capturing the complexities of neural criticality.

To quantify the hierarchical arrangement and irregularity (stability) between frequency oscillations, researchers have turned to entropic methods, particularly spectral entropy [29]. Spectral entropy, derived from Shannon entropy [29, 46], uses the spectral intensity of each frequency oscillation within a power spectrum of brain activity as probabilities for entropy calculation. A high spectral entropy value indicates a uniform distribution of spectral intensity across frequencies, while a low value reflects a power spectrum concentrated within a singular frequency bin [1]. Although spectral entropy has been used to explore the mechanistic characteristics of neural activation and firing through dominant frequency bands, it fails to capture irregularities and hierarchical arrangements in frequency band activity.

To address temporal irregularity, studies have employed other entropic methods, such as sample entropy [9]. While sample entropy captures temporal irregularity, it forfeits the ability to characterize the spectral features of hierarchical arrangements between neural oscillations. This disconnect creates a gap in understanding transitional neural activation associated with criticality, as current entropic methods fail to simultaneously capture both the spectral and temporal arrangements of neural oscillations. Consequently, the hierarchical arrangements and stability of neural communication, along with its underlying processes (e.g., points of criticality), remain unexplored. There is currently no metric capable of fully capturing the interactions of neural oscillations and how they arrange over time, leaving critical dynamics of neural activation and cognitive processes insufficiently understood.

## 1.3. Insights

As previously discussed, classical wavelet filter banks lack the spectral resolution necessary for accurate one-to-one intensity analysis of EEG band activity, resulting in redundant spectral information across multiple wavelet scales. Traditional wavelet methods, such as the continuous wavelet transform, are primarily designed to localize frequency content and transient behaviors within a signal. However, they do not inherently provide the direct intensity of the signal, which is critical for the interpretability of EEG data. This limitation hinders the ability to statistically compare the intensity of EEG frequency bands (neural oscillatory groups).

Optimized filter bank designs leveraging wavelet concepts have been developed to overcome these limitations, enabling one-to-one representation of intensity for EEG frequency oscillations (i.e., delta, theta, alpha, beta, and gamma) over time [24]. This unique design allows for precise temporal analysis of the dynamics, intensity, and interactions of neuronal oscillations. These interactions may reflect synchronized neural activity, facilitating communication between oscillatory groups essential for cognitive processes. By analyzing critical points during cognitive processes, such filter banks could reveal enriched neural information vital for understanding cognitive mechanisms, task detection, and impaired brain function during cognitive tasks.

To investigate the mechanisms underlying neural criticality and communication, we utilize a hypoxia dataset in which participants perform cognitive tasks. Hypoxia, a condition characterized by insufficient oxygen supply at the tissue level, disrupts neural communication and firing due to ATP depletion [31, 39, 24]. This disruption impairs brain function during cognitive tasks. Examining the effects of hypoxia on neural communication and criticality activation provides valuable insights into the dynamics of neural communication and its relationship to cognitive states and impairments.

While spectral intensity dynamics offer significant insights into cognitive states [46, 37], current metrics are limited to static snapshots of the frequency spectrum (as illustrated in the central plot of Figure 2). This static approach overlooks the unique signatures of neural communication between oscillatory groups during criticality events, which may vary depending on the specific cognitive process or state. These variations are reflected in the shifting dominance of neural oscillations over time, as depicted in the surrounding plots of Figure 2. Capturing the temporal hierarchical arrangement of these oscillations is crucial for characterizing neural communication patterns and understanding transitions of criticality during cognitive processes.

## 2. Methods: Data Collection and Breakdown of Cognitive Tasks

### 2.1. Hypoxia Dataset

The study at NASA Langley Research Center (LaRC) investigated the impact of mild hypoxia on cognitive performance in aircraft pilots. With approval from the Institutional Review Board, 49 participants (41 males, 8 females; average age 43.3 years) gave informed consent. Before hypoxia was induced, all participants underwent screening for neurological and cardiovascular conditions. Each participant also had current hypoxia training certifications for normobaric hypoxia and some flight experience. Simulated altitudes of sea level (21% *O*_2_) and 15,000 feet (11.2% *O*_2_) were induced using an Environics, Inc. Reduced Oxygen Breathing Device (ROBD-2). During both conditions, participants completed 10-minute sessions of the Multi-Attribute Task Battery (MATB-II) [38] to assess cognitive performance. EEG data were collected using a 16-lead system, selected to minimize interference from the ROBD-2 oxygen mask (see Figure 3). Since participants were required to breathe through the mask, its backstrap could obstruct certain electrode placements. To ensure high-quality recordings while accommodating the mask, we positioned electrodes to avoid interference while maintaining as close an alignment as possible with the standard 10-20 system.

**Figure 3:**
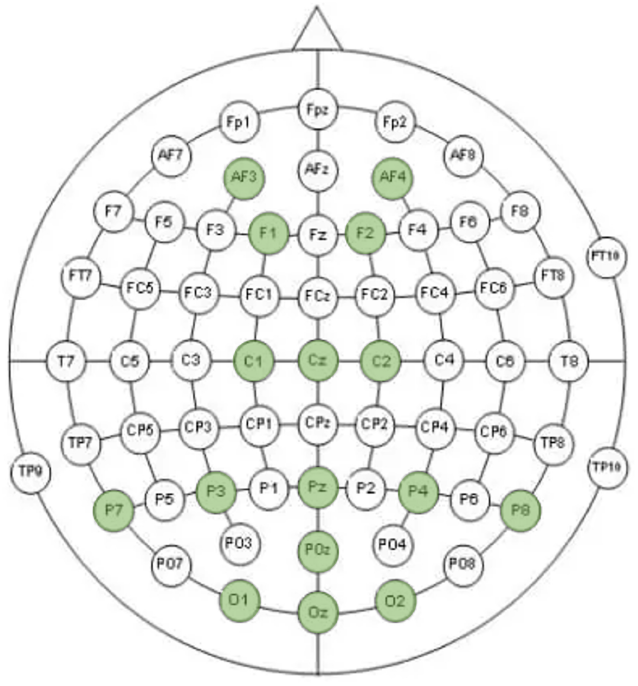
A visual representation of the EEG electrode placements used in this study, highlighted in green.

### 2.2. Explanation of MATB-II Operation

The NASA MATB-II [38] presents a single interface (Figure 4) with four task components: Tracking, Communications, Resource Management, and Systems Monitoring. For this study, analysis focused on the Tracking, Communications, and Resource Management tasks. All data were collected during the manual tracking mode, excluding the automatic tracking option. In the tracking task, participants used a joystick to keep a crosshair centered within a square on the display (Figure 4). The communications task simulated radio communication, requiring participants to listen for their assigned call sign. If the call sign was mentioned, they adjusted the specified radio and frequency channel. If not, participants ignored the audio and maintained the current settings. Adjustments were made using either a mouse or a keyboard.

**Figure 4:**
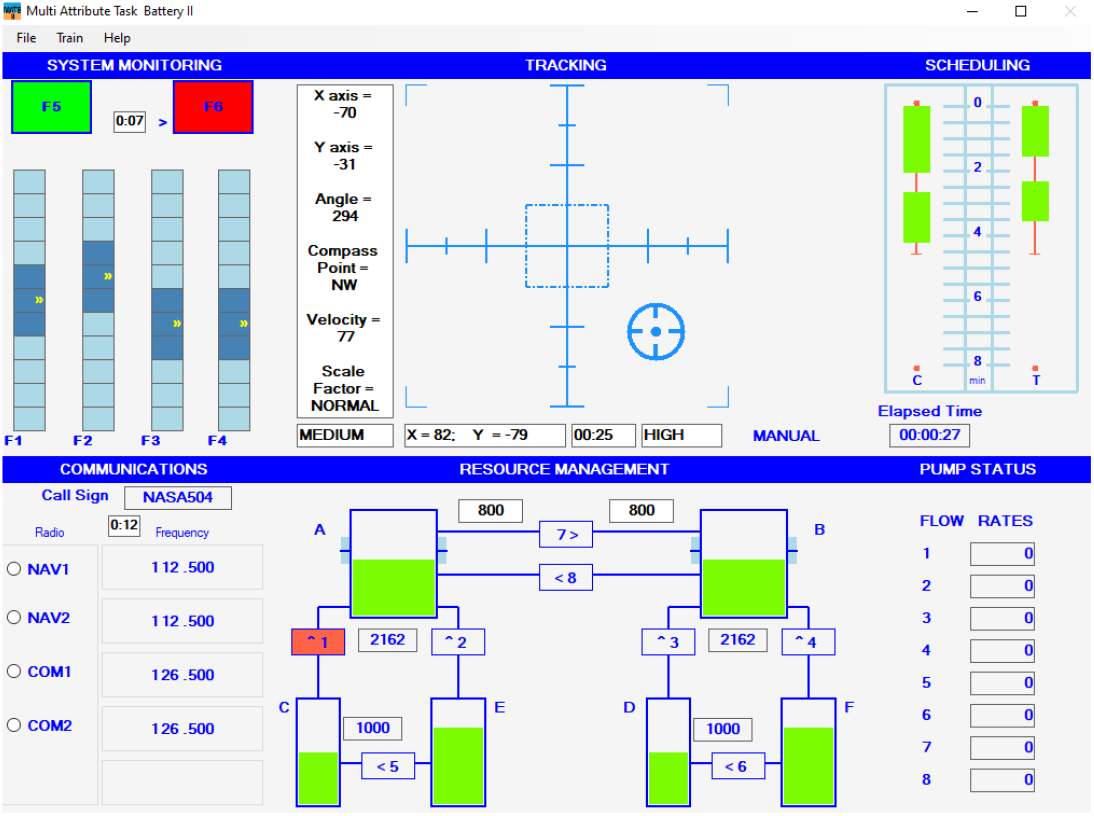
The NASA MATB-II layout utilized in this study. Our analysis focuses exclusively on responses related to the Tracking, Communications, and Resource Management tasks.

### 2.3. MATB Output and Ground Truth Assignment

The raw MATB output files were analyzed line by line to extract action statements and timestamps, which were then used to define the endpoints of distinct cognitive task states. This approach is novel, as MATB files have not previously been examined in this way to systematically define cognitive task states. By structuring MATB output into distinct cognitive phases, we introduce an innovative method for extracting insights into multitasking behavior and neural activity.

For the communication task, system-generated remarks were used to align key actions—such as task initiation, user responses, and task completion—with specific cognitive states, as shown in Figure 5. Based on these remarks, we identified five distinct cognitive states:

**Figure 5:**
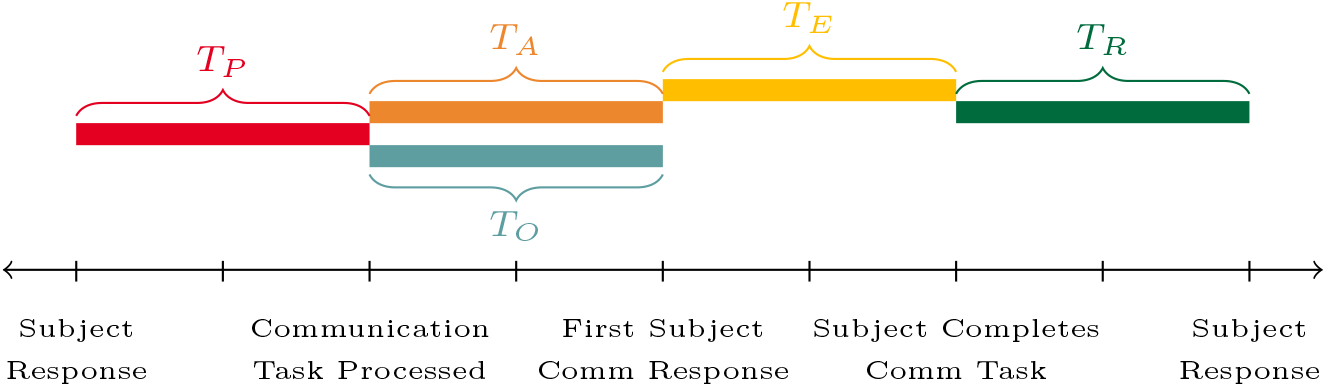
This figure provides a breakdown of the variables derived from the deconstruction of a MATB task. The variables *T*_*P*_, *T*_*A*_, *T*_*E*_, *T*_*R*_, and *T*_*O*_ correspond to defined task states, representing different time windows within a task: task precursor, audio task interruption, task execution, task recovery, and other task, respectively. The events or actions that define the endpoints of these states are depicted along the x-axis of the timeline.

- **Task Precursor (***T*_*P*_) – The period before the communication task begins. This state captures early neural interactions before the introduction of a secondary task, setting the stage for multitasking.
- **Audio Task Interruption (***T*_*A*_) – The moment when the participant first hears and responds to an audio cue, marking the onset of multitasking. It specifically occurs when the participant’s assigned call sign is mentioned, requiring them to adjust the radio frequency. This state reflects synchronized neural activity and auditory processing, including the participant’s auditory brainstem response.
- **Task Execution (***T*_*E*_) – The active phase of performing the communication task while simultaneously managing the tracking task. This state captures neuronal firing patterns and inter-group communication associated with task-switching and motor-related cognitive function.
- **Task Recovery (***T*_*R*_) – The transition period following task completion but before responding to the next event. This state reflects the brain’s shift from a high-workload multitasking condition to a lower workload demand, potentially revealing changes in neuronal communication patterns.
- **Task Other (***T*_*O*_) – The state in which participants were exposed to an audio cue but their assigned call sign was not mentioned. In this scenario, they were required to ignore the cue and maintain the current radio frequency settings. This state represents a passive listening condition where participants remained attentive but did not need to execute an action, providing a contrast to the multitasking nature of *T*_*A*_.

## 3. Methods: Proposed Algorithms to Characterize Neuronal Dynamics

This section outlines the evaluation of neuronal oscillatory bands over time, as motivated by Table 1 and Figure 2. The EEG waveform is decomposed using an optimized filter bank approach [24, 26], which forms the basis of the various proposed algorithms. This allows us to compare the different types of information (e.g., temporal, spectral, variance of oscillatory groups, hierarchical arrangements) required to decode the neural oscillations.

### 3.1. Preprocessing with Established Filter Bank for Spectral Intensity Extraction

We deployed the proposed filter, as detailed in [24, 26], to extract each EEG frequency band and analyze the continuous time-frequency intensity dynamics of EEG signals. This design was chosen because it ensures accurate extraction of each EEG frequency band while maintaining appropriate filter bank plateau values. To meet these and other constraints, a flattened Gaussian basis function 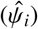 was employed [6]. The basis function that creates each *i*^*th*^ filter of the filter bank is defined as

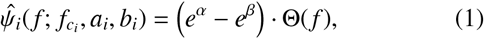

where 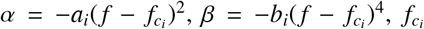 is the center frequency, *a*_*i*_ and *b*_*i*_ are the tuning parameters, and Θ(*f*) is the Heaviside function to constrain the filter to only positive frequencies [26]. The basis function parameters were optimized to ensure the filter bank uniformly represented all EEG frequencies while maintaining reasonable separability between the frequency bands. The optimized parameters are summarized in Table 2, including the center frequencies 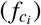, weights (*a*_*i*_ and *b*_*i*_), and lower and upper cutoff frequencies 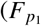 and 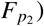 for each filter. Once designed in the frequency domain, the filters are transformed into the time domain using the Inverse Fourier Transform, followed by a circular shift [24], resulting in 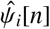. The EEG signal, *x*_*s*_[*n*] is then convolved with 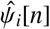 producing *ρ*_*i*_[*n*], where each *i* output is associated with its respective 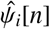. This spectral output intensity, *ρ*_*i*_[*n*], is then convolved with *G*_*f*_ [*n*], a Gaussian Smoothing Filter,

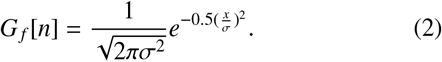

where 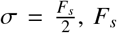 is the sampling frequency and 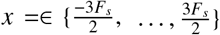. This produces a smoothed spectral output intensity, 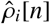, where the details of the full process are described in [24].

**Table 2:**
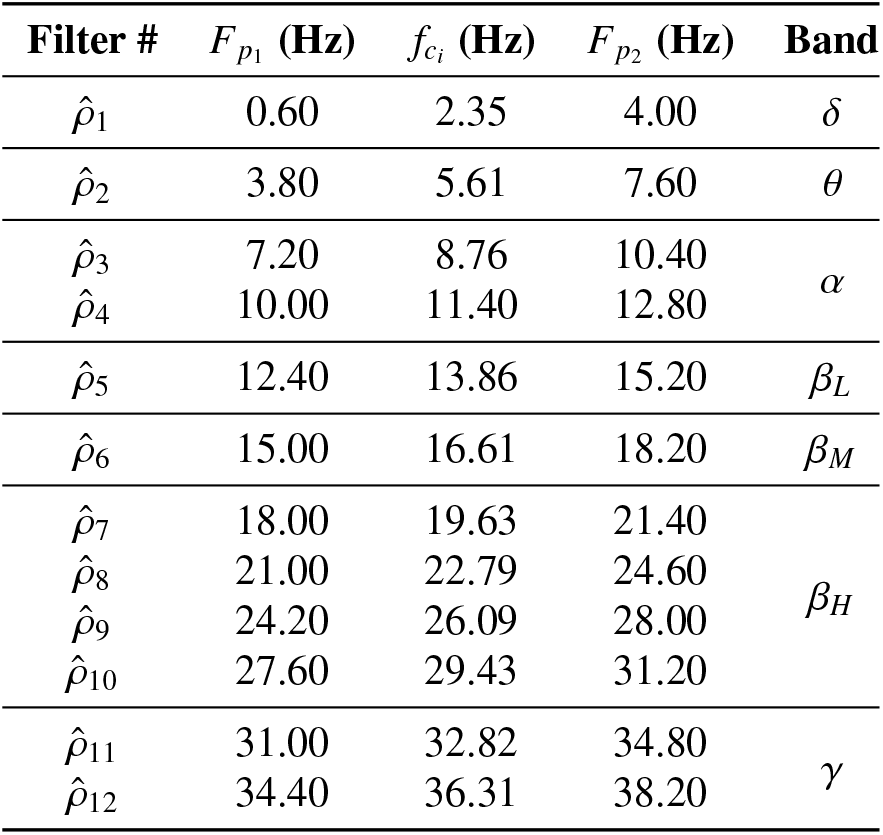
Filter Bank Parameters.

**Table 3:**
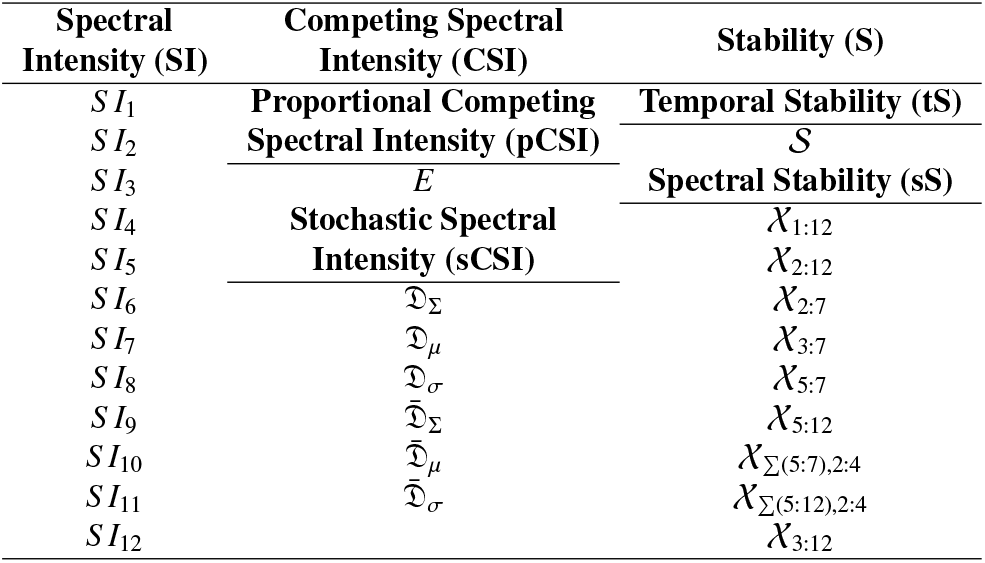
Notation for EEG Features Extracted.

### 3.2. Proposed Algorithms for EEG Dynamics Analysis

This subsection presents four main conceptual approaches for decomposing and evaluating the dynamics of hierarchical arrangements within a critical event generated by these neuronal oscillatory bands: 1) Spectral Intensity (SI) in Section 3.2.1; 2) Variations of Competing Spectral Intensity Measurements (CSI) in Section 3.2.2; 3) Spectral Stability (SS) in Section 3.2.3; and Spectral Intensity Stability (SIS) is Section 3.2.4.

#### 3.2.1. Filter Bank Bands: “Spectral Intensity (SI)”

Spectral intensity is a standard metric of EEG interpretation, we use this first measure as the baseline or control to compare to the other methods. Utilizing the foundational EEG filter bank, we can calculate the total intensity for each spectral band and for each task (e.g., *T*_*p*_). This is accomplished by first defining the window, *τ*, for each task from the MATB file, which is delineated by the start time, *n*_*s*_, and a completion time, *n*_*c*_ of the task, described in Subsection 2.3. The total spectral intensity is calculated by,

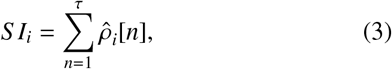

where *i* is the filter.

#### 3.2.2. Competing Spectral Intensity (CSI)

The Engagement Index (EI) was introduced by Pope et al. as a psychophysiological measure that indicates the level of engagement of the user in a task based on the proportional power of the alpha, beta, and theta EEG bands [32]. The underlying principle is based on the frequency band interpretation: the beta EEG frequency band is linked to concentration, active thinking, and arousal. Thus, beta is set proportionally at the numerator, while the EEG frequency bands of delta and theta, which are linked to sleep and inhibition, are set at the denominator. This allows us to compare the competing nature between EEG frequency bands through proportional changes. This work has been extensively used in literature to monitor human engagement during tasks, which has been related to workload as task load increases [33].

In this section, we introduce the classical approach of utilizing the EI, *E*, which allows us to compare the proportional nature of frequency bands but fails to provide information about how spectral intensity dynamics compete and change over time. We then propose a new dynamic method for the EI that allows us to capture the stochastic nature of the EEG’s spectral intensity dynamics over time in a more granular way, denoted as 𝒟 ^*k, j*^[*n*]. 𝒟 ^*k, j*^[*n*] is explored over two different ranges of beta and then described using summary statistics of sum (Σ), mean (*µ*), and standard deviation (*σ*).

##### Proportional CSI (pCSI)

The Classical EI was the first algorithm introduced by Pope. It is calculated using the proportion of beta over the sum of alpha and theta waves [32], where we refer to this as Proportional CSI (pCSI). The EI defined over a window of time *τ* is defined as,

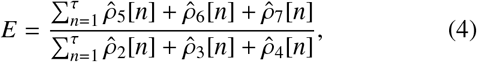

where 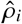 individual over a time window. However, when the time window becomes too large, multiple cognitive states may occur and attention may vary. Therefore, this approach is limited due to a lack of temporal information that specifies how engaged an individual was at a specific point in time.

##### Stochastic CSI (sCSI)

Expanding the EI analysis by examining how engagement dynamically alters over time, we utilize the mean and standard deviation to capture the stochastic nature of these oscillatory bands, where we refer to this as Stochastic CSI (sCSI). As mentioned previously, the original EI method is a summation of the intensity over a window of time *τ*. As a result, it does not provide the temporal dynamics of the EEG frequency bands within the window of time *τ*, only the overall engagement. To address this, we propose a dynamic engagement approach to the EI, defined as 𝒟 ^*k, j*^[*n*], which is an instantaneous measurement of the EEG intensity found within the filter bank outputs, 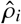. For each point *n* in time, the EI is quantified as follows:

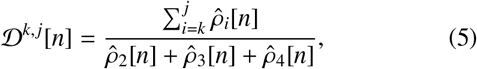

where k and j are the lower limit and upper limit intensity filter banks, respectively. This change in the calculation provides an instantaneous overview of EI changes over time within the window *τ*.

The definition of 𝒟 _*k j*_ is expanded to characterize the dynamics of the EI over a window of time *τ*. The sum, mean, and standard deviation of the EI over this window of time *τ* are described as:

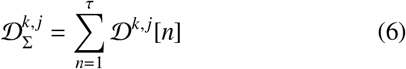

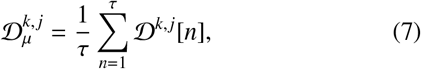

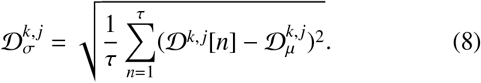

Both 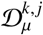 and 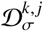 are similar to the original engagement index *E* in that they are static measurements of task engagement within a specified window of time. However, 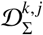 differs because the summation provides a more dynamic measurement of task engagement, capturing the total task engagement seen within a window of time. Figure 6 depicts an example of how CSI is calculated. The proposed method explores two different frequency ranges for the numerator, but algorithmically, various hierarchical arrangements should be further investigated. Our first frequency bands utilized within the numerator are the original frequency ranges applied for the classical EI, which are 13-22 Hz and denoted as 𝒟 ^5,7^. An extended frequency range is also explored, which encompasses the low beta, mid beta, high beta, low gamma, and high gamma spectral intensity bands. This second hierarchical arrangement extends the frequency range from 13-22 Hz to the range of 13-38 Hz. This extended range is defined as 𝒟 D^5,12^.

**Figure 6:**
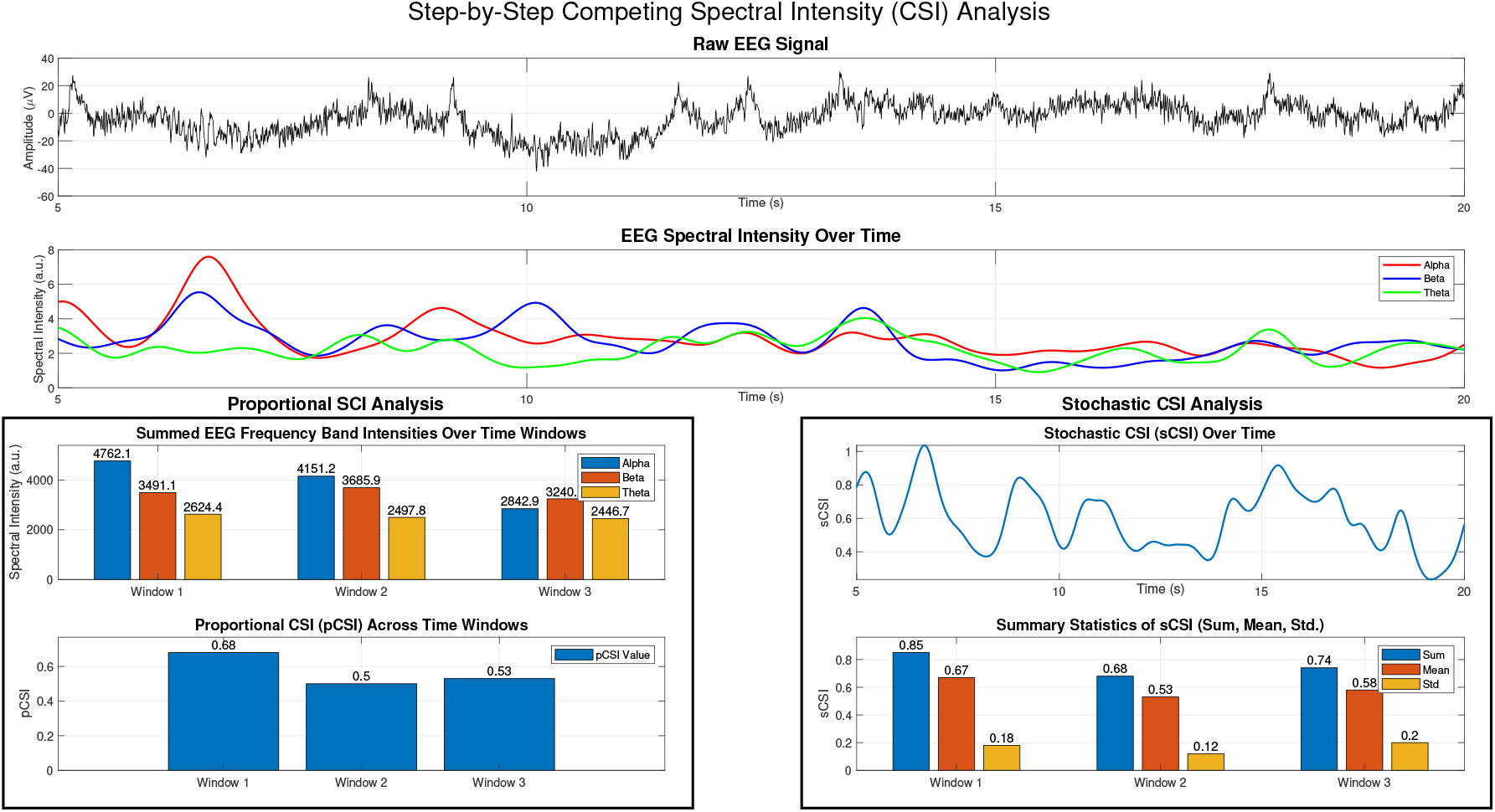
Step-by-step visualization of the Competing Spectral Intensity (CSI) analysis. The top panel shows the raw EEG signal over time. The middle panel displays the spectral intensity of the alpha (red), beta (blue), and theta (green) frequency bands, which are used to compute CSI. The middle-left panel presents the summed spectral intensity of these bands over three non-overlapping 5-second time windows, highlighting their proportional contributions. The bottom-left panel shows the Proportional CSI (pCSI) across each time window, representing a static measure of engagement. The middle-right panel depicts the Stochastic CSI (sCSI) over time, capturing the dynamic fluctuations in engagement. The bottom-right panel summarizes the sum, mean, and standard deviation of sCSI across each 5-second time window. To ensure better visualization, the sum of sCSI has been scaled down by a factor of 1000 to match the scale of the mean and standard deviation.

#### 3.2.3. Oscillations Group Stability “Spectral Stability (SS)”

Spectral Stability (SS) quantifies how stable or predictable the hierarchical arrangements of neuronal oscillations are over time. We evaluate SS using an entropy-based metric called Temporal Cross-Band Entropy (TCBE), which measures the disorder or unpredictability in the rank order of spectral intensities across different EEG frequency bands.

Building on the foundational EEG filter bank, specific hierarchical arrangements can be identified through rank-order patterns at each time instance, enabling evaluation of the unique dynamics of oscillatory group arrangements. This algorithm is versatile and can be applied to time series data of varying lengths. Instead of treating the signal as a single one-dimensional sequence, it is analyzed as a matrix, where each row corresponds to an output of the filter bank, and each column represents a time instance. For the proposed filter bank, we have *M* = 12 filters and a given time series*x*_*s*_[*n*], we produce *M* intensity sequences, 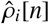, where ∀*i* = 1, …, *M* and ∀*n* = 1, …, *N*. This allows us to form an intensity matrix,

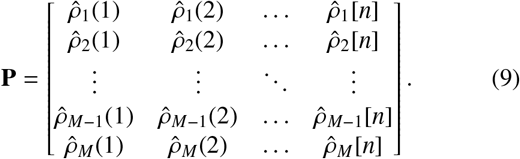

Similar to how all other entropy measurements examine a vector sequence with a defined template length [21] or an embedded dimension [49], our embedded dimension is defined by the number of filters in the filter bank, *M*. Thus we examine *N* column vectors composed from **P**, which consist of *M* subsequent intensity values. Each column vector,

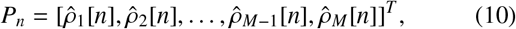

has a specific rank order pattern which is defined as a unique hierarchical arrangement (permutation), *π*_*C*_[*n*] ≜ [*r*_1_, *r*_2_, …, *r*_*M*_] of the form (1, 2, …, *M*), which fulfills

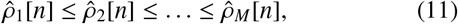

where *C* ∈ {1 … *M*!} and *π*_*C*_ equal of total number of occurrences of that hierarchical arrangement, *C*. For clarification, a numerical example is provided, starting with the intensity matrix,

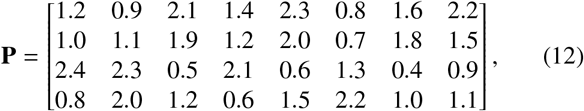

where the embedded dimension is *M* = 4 and the length of the time series is *N* = 8. In this case, the matrix **P** produces multiple unique hierarchical arrangements. For example:

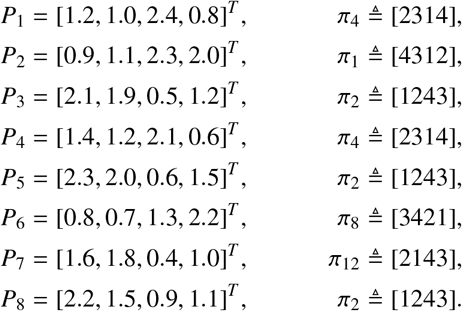

Thus the, total number of unique hierarchical arrangements that can be achieved is *M*! = 4! = 24, and *π*_*C*_ equals the number of occurrences for each unique hierarchical arrangement: *π*_1_ = 1, *π*_2_ = 3, *π*_4_ = 2, *π*_8_ = 1, *π*_12_ = 1, and others= 0.

From these unique hierarchical arrangements, we evaluate spectral stability using entropy, a measure of disorder or predictability in time series data. Traditional entropy methods are often used in nonlinear systems to assess complexity or stability but are primarily designed for time-domain analysis rather than spectral information [14, 42, 24]. To address this, we introduce Temporal Cross-Band Entropy (TCBE), an entropy-based method tailored to quantify spectral stability by analyzing how the hierarchical arrangements of neuronal oscillations evolve over time. Unlike conventional entropy measures, TCBE accounts for both spectral and temporal dynamics while disregarding absolute intensity values. TCBE evaluates how spectral bands interact by tracking their rank order over time. When the rank order remains stable, entropy is low, indicating high spectral stability. Conversely, frequent changes in rank order result in higher entropy, reflecting increased variability in spectral organization. Unlike conventional entropy measures, TCBE operates in a two-dimensional space and does not require a predefined template length [49, 25]. The dynamics of spectral intensity rank changes analyzed through TCBE are illustrated in Figure 7.

**Figure 7:**
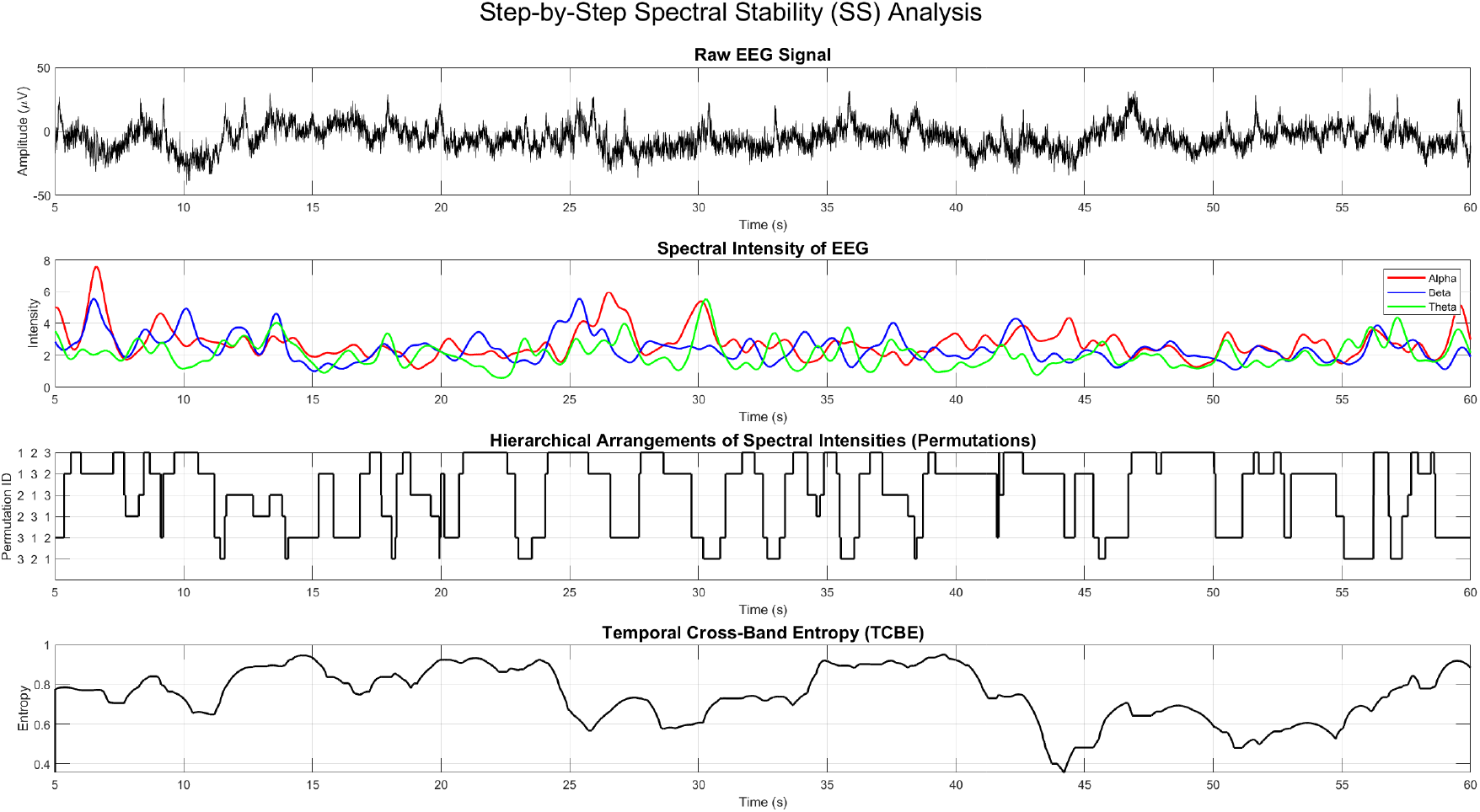
Step-by-Step Illustration of Spectral Stability (SS) Analysis. This figure demonstrates the calculation of spectral stability by tracking the hierarchical ordering of EEG frequency bands over time. The top panel presents the raw EEG signal. The second panel shows the intensity of the alpha, beta, and theta bands, which form the basis of the stability analysis. The third panel displays the hierarchical rank order of these frequency bands, where each unique permutation is assigned a numerical label and plotted as a step function. The bottom panel shows TCBE, which quantifies spectral stability by measuring the variability of rank-order transitions over a time window. The zero values at the beginning of the TCBE signal illustrate that TCBE is computed over a window of time (5 seconds in this example), meaning entropy values cannot be determined for the first few seconds of data. Higher TCBE values indicate greater instability in spectral ordering, while lower values reflect more stable hierarchical arrangements of EEG frequency bands.

The TCBE metric, X, is computed as:

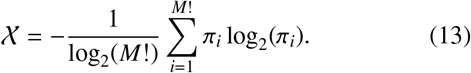

The notation χ _*a*:*b*_ was coined to describe the TCBE across the range of filters from *a* to *b*, with *M* the total number of filters. We also coin the notation χ _(*c*:*d*),*a*:*b*_ where in addition to filters *a* to *b*, we include an additional composite filter of the summation of filters *c* to *d*. In either case, the intensity matrix **P** would be manipulated to account for this new range. Thus, each column vector, *P*_*n*_, of the intensity matrix would be defined as,

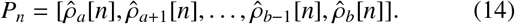

For example, when we declare χ_5:12_, it means we applied Equation 13 to the unique hierarchical arrangements found in the intensity matrix

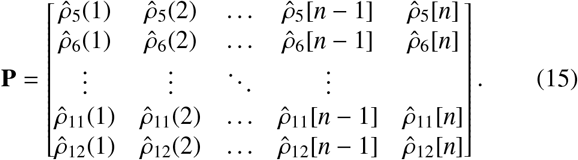

#### 3.2.4. TCBE x SI (“Spectral Intensity Stability (SIS))”

Spectral Intensity Stability (SIS) captures the relationship between spectral intensity and spectral stability, providing insights into how the strength and arrangement of oscillatory bands evolve over time. SIS is computed as a weighted interaction between TCBE and spectral intensity:

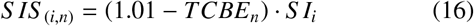

where *S I*_*i*_ represents the spectral intensity of the *i*^*th*^ frequency band, and *TCBE*_*n*_ quantifies the spectral stability at time *n*. The factor 1.01 is introduced because TCBE is bounded between 0 and 1, ensuring that SIS does not become excessively large when TCBE values are very small, maintaining a more balanced measure of spectral intensity stability.

One of the key strengths of SIS is its high temporal resolution. Since EEG was sampled at 256 Hz in this study, each time step *n* corresponds to approximately 3.91 milliseconds. This allows SIS to capture rapid fluctuations in neural oscillatory hierarchies that may be missed by traditional sliding-window spectral analysis. By updating TCBE and spectral intensity at each sample, SIS offers a fine-grained view of neural dynamics, making it especially useful for detecting short-lived critical transitions in cognitive state.

To manage the complexity introduced by the numerous possible hierarchical arrangements, we focus on the most predictive spectral stability features, specifically χ_5:12_, which corresponds to beta and gamma bands, frequencies closely associated with cognitive processes. However, SIS can be extended to various frequency combinations, making it a versatile tool for future research on neuronal dynamics.

## 4. Methods: Evaluating EEG Feature Families for Cognitive Task Prediction

### 4.1. Controlled Feature Evaluation for Explainable Cognitive State Detection

To systematically evaluate the discriminative power of our proposed EEG-based algorithmic strategies, we conducted a controlled feature evaluation, analogous to an ablation study. Each algorithmic strategy was assessed under identical experimental conditions to ensure a fair and interpretable comparison. Specifically, we constructed four independent models, each using features exclusively from one of the proposed feature families (SI, CSI, SS, or SIS). All models shared the same machine learning architecture based on the NAPS framework and were restricted to the same number of features (N) drawn from their respective families. Additionally, all experiments used identical dataset splits, class label definitions, and evaluation protocols. This controlled setup allowed us to isolate the contribution of each feature family and quantify its predictive utility for cognitive state classification. Model performance was evaluated using the F1-Score and the Matthews Correlation Coefficient (MCC)[13, 23, 45]. A visual summary of this evaluation framework is provided in Figure 8.

**Figure 8:**
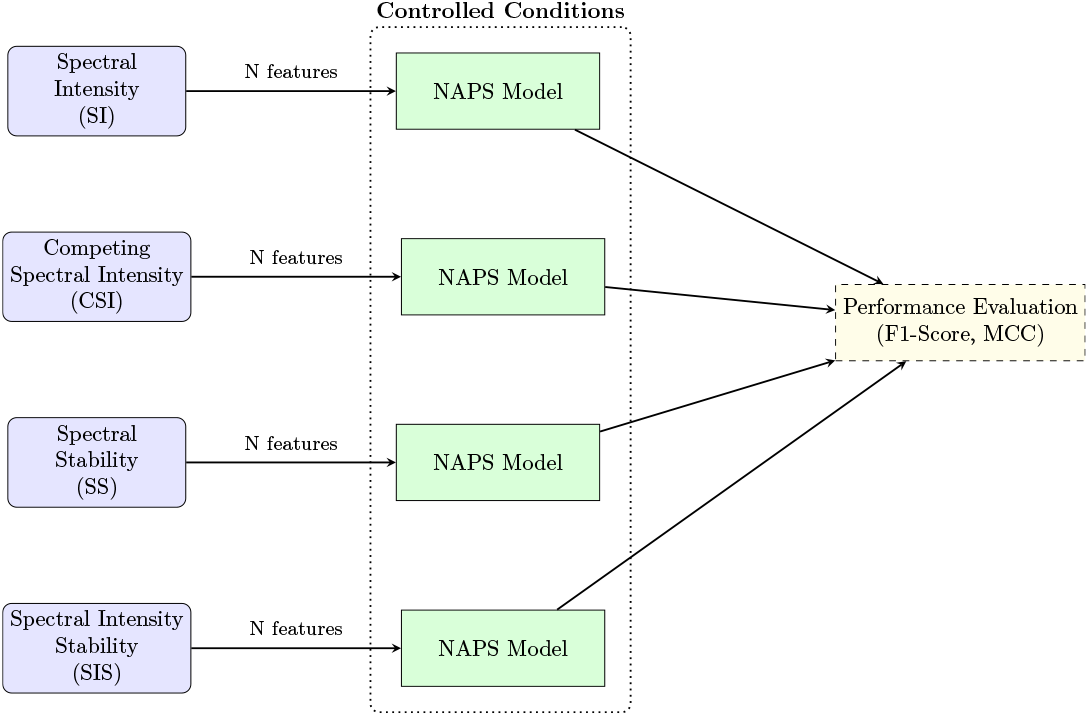
Controlled feature evaluation framework. Each feature family (SI, CSI, SS, SIS) was evaluated using an identical NAPS-based model with the same number of features, dataset splits, and evaluation protocols.

The classification problem was based on task-related cognitive states derived from MATB-II log file decomposition (detailed in Section 2.3), resulting in five distinct classes: Task Precursor (*T*_*P*_, 411 samples), Task Interruption (*T*_*A*_, 411), Task Execution (*T*_*E*_, 411), Task Recovery (*T*_*R*_, 411), and Task Other (*T*_*O*_, 460). To incorporate the effects of cognitive impairment, each class was further split into hypoxia (H) and non-hypoxia (NH) conditions, forming a ten-class problem. The hypoxia conditions included 212 samples each for 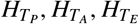, and 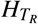, with 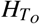 comprising 230 samples. The non-hypoxia counterparts had 199 samples each for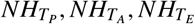, and 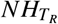, and 230 samples for 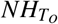.

### 4.2. NAPS-Augmented Machine Learning Framework

The machine learning model used in this study is the NAPS Augmented ML Framework, originally developed to support physiological data analysis such as EEG [27]. NAPS was chosen because it is designed for tabular feature data and uses interpretable decision tree models, making it well-suited for identifying which features drive classification performance. It includes two components particularly useful for this work: augmented response variable mapping and a sensor model paradigm, both of which improve the ability to model subtle variations across cognitive states. While we did not use the full NAPS pipeline (e.g., fusion and uncertainty weighting), we implemented its core structure: the all-versus-one augmented response mapping and the sensor model architecture. These components were used to generate hierarchical combinations of response variables and enable robust modeling across subject-specific data subsets. After applying the sensor model paradigm, we trained models using Bootstrap Aggregated Decision Trees (Tree-Bagger in MATLAB) [23], which aggregate multiple decision trees created via Breiman’s Random Forest algorithm [8]. This ensemble approach improves generalizability and reduces overfitting, especially in high-variance physiological data. Voting across the sensor models was used to derive final predictions. To handle class imbalance, especially in the binary augmented models, the Synthetic Minority Oversampling Technique (SMOTE) was applied during training. An example of how NAPS generates augmented binary models from a five-class structure is shown in Table 4. To answer RQ2, feature inputs were restricted to a single algorithmic strategy per model to evaluate which feature family (SI, CSI, SS, or SIS) best captured task-related cognitive state transitions. Each sensor model used a fixed number of features from one family, and the final classification was based on the 16 best-performing models with the least associated uncertainty.

**Table 4:** Example of a 5 Class Augmented Response.

| Combination Variations | Class / Proposition Assignments | Number of Classes |
| --- | --- | --- |
| $A_0$ | $\{T_P\} \{T_A\} \{T_O\} \{T_E\} \{T_R\}$ | 5 |
| $A_1$ | $\{T_P\} \{T_A, T_O, T_E, T_R\}$ | 2 |
| $A_2$ | $\{T_A\} \{T_P, T_O, T_E, T_R\}$ | 2 |
| $A_3$ | $\{T_O\} \{T_P, T_A, T_R, T_E\}$ | 2 |
| $A_4$ | $\{T_E\} \{T_P, T_A, T_O, T_R\}$ | 2 |
| $A_5$ | $\{T_R\} \{T_P, T_A, T_O, T_E\}$ | 2 |

## 5 Results

To determine which neuronal firing dynamics contain the most information for cognitive state detection, we examine the four proposed families of features to evaluate the most discriminative for the detection of cognitive task states and impairment (i.e., hypoxia). These families of features all use the same foundational spectral decomposition method but process the decoding of the activation and interaction of neural oscillatory bands differently. This decoding is conceptually focused on evaluating how the intensity dynamics of neural frequency-based oscillations alter or are arranged hierarchically over time. These nascent methodological approaches provide a critical foundation for understanding underlying dynamics and processes of criticality concerning cognition that have been neglected. This evidence supporting the specific mechanics of neuronal firing is through the information extracted in order to discriminate cognitive states which are substantiated by assessing the predictive performance of the ML model (discussed in Subsection 4.2) through a comparison of models using various contextual features that characterize EEG dynamics in four primary dimensions: 1) Spectral Intensity, 2) Spectral Stability, 3) Competing Spectral Intensity, and 4) Spectral Intensity Stability. The optimal predictive model identifies which of these features offers the most distinguishing and informative EEG dynamics for accurately predicting cognitive processes in task classification. In our reporting of the ML’s performance, we provide two summary metrics: the MCC and F1 scores. In addition, four supporting metrics provide further insight into the performance and errors: false positives (FP), false negatives (FN), true positives (TP), and true negatives (TN). The FPs, also known as the false alarm, occur when the class is classified as positive even when it is not. The FNs, also known as the miss classification, occur when the classification model fails to detect a true positive.

To ensure the validity of cognitive state classification, ground truth labels were established based on a structured decomposition of MATB output logs (see Subsection 2.3). The MATB system provides precise event markers, including task initiation, user responses, and task completion timestamps, which were used to define transitions between cognitive states: Task Precursor (*T*_*P*_), Audio Task Interruption (*T*_*A*_), Task Execution (*T*_*E*_), Task Recovery (*T*_*R*_), and Task Other (*T*_*O*_). These transitions were objectively determined using system-generated logs, ensuring that state definitions were not based on subjective criteria. Additionally, timestamps from these task events were aligned with EEG data to verify that neural activity patterns corresponded to distinct cognitive states.

### 5.1. RQ1: Decoding Neuronal Dynamics Related to Task Classification

Our first research question is “Do unique dynamics within these task events that elicit criticality have unique dynamics, potentially establishing a “fingerprint” for cognitive processes?”. We first present the predictive model’s performance for the merge cohorts of the hypoxia and non-hypoxia datasets for the 5-class problem across each algorithmic strategy.

The summary metrics of MMC and F1-Score across algorithmic strategies for the 5-class model are depicted in Figure 9, where the traditional *SI algorithm* achieves an overall MCC of 0.52 and is the lowest performing model presented within the set. The *CSI algorithmic strategy* exhibits an MCC of 0.63 for the 5-class classification task, providing a predictive power increase of 21.15 % compared to the SI. The SS algorithm achieves an MCC of 0.54 with a 3.85% increase compared to the SI algorithm and a 16.67% decrease compared to the CSI algorithm. However, most importantly, the *SIS algorithmic strategy*, shows the best performance with an MCC of 0.68 with an increase of 7.93% when compared to the CSI, a 20.58% increase when compared to SS and 23.52% increase compared to the traditional SI algorithm.

**Figure 9:**
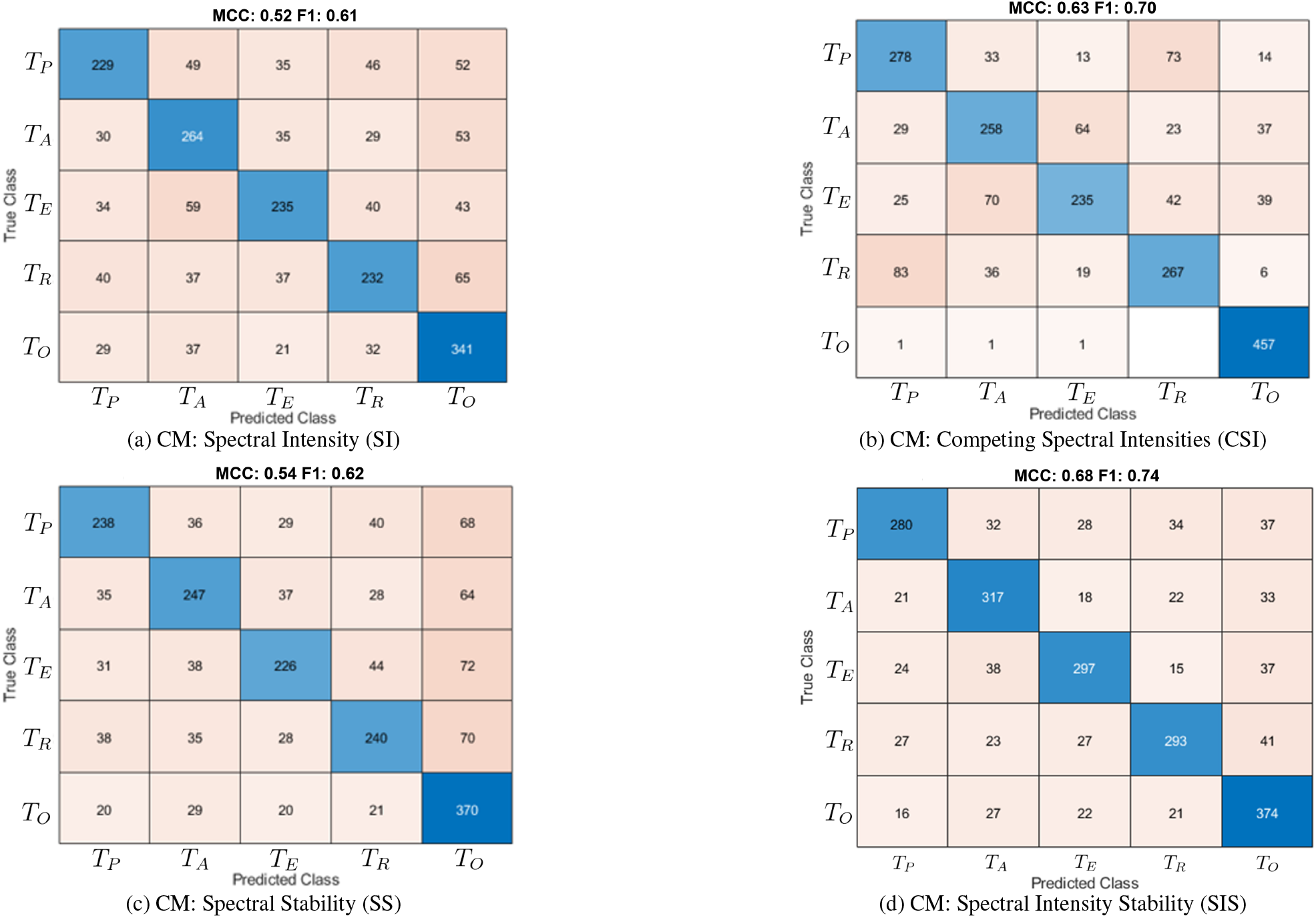
The four confusion matrices (CM) compare the predictive performance of the following methods: (1) classical spectral intensity (SI) analysis, which provides intensity information without incorporating temporal dynamics; (2) the proposed dynamic engagement index, which measures the stochastic properties of spectral intensity dynamics; (3) the proposed TCBE, which captures spectral temporal dynamics while excluding intensity information; and (4) the interaction of TCBE with spectral intensity, representing temporal spectral intensity dynamics.

Table 5 exhibits a breakdown of each algorithm’s performance using the true positives (TP), true negatives (TN), false positives (FP), and true negatives (TN) across the five classes. This table allows us to evaluate if a unique algorithmic strategy can discriminate across a specific cognitive task and how it performs overall. It is worth noting that SIS dominating performs across all tasks and for all performance metrics (e.g., TP, TN) except for *T*_*O*_. The CSI algorithm best predicts the *T*_0_, with an increase of 18.16% of TPs compared to the second-best algorithm, SIS. Although the worst-performing algorithm for predicting TPs is SI, SS performs only slightly better.

**Table 5:** This table compares all induced tasks across the four algorithmic strategies, presenting the number of True Positives, False Negatives, False Positives, and True Negatives. The highlighted cells provide a visual comparison of performance: dark green indicates the best performance, light green denotes the second-best performer, orange marks the second-to-worst performer, and red highlights the worst performer. This coloring scheme aids in identifying which algorithmic strategy is most favorable for specific tasks.

|  | True Positive |  |  |  | False Negative |  |  |  |
| --- | --- | --- | --- | --- | --- | --- | --- | --- |
|  | SI | CSI | SS | SIS | SI | CSI | SS | SIS |
| $T_P$ | 229 | 278 | 238 | 280 | 182 | 133 | 173 | 131 |
| $T_A$ | 264 | 258 | 247 | 317 | 147 | 153 | 164 | 94 |
| $T_E$ | 235 | 235 | 226 | 297 | 176 | 176 | 185 | 114 |
| $T_R$ | 232 | 267 | 240 | 293 | 179 | 144 | 171 | 118 |
| $T_O$ | 341 | 457 | 370 | 374 | 119 | 3 | 90 | 86 |

Table 5: This table compares all induced tasks across the four algorithmic strategies, presenting the number of True Positives, False Negatives, False Positives, and True Negatives.
|  | False Positive |  |  |  | True Negative |  |  |  |
| --- | --- | --- | --- | --- | --- | --- | --- | --- |
|  | SI | CSI | SS | SIS | SI | CSI | SS | SIS |
| $T_P$ | 133 | 138 | 124 | 88 | 1560 | 1555 | 1569 | 1605 |
| $T_A$ | 182 | 140 | 138 | 120 | 1511 | 1553 | 1555 | 1573 |
| $T_E$ | 128 | 97 | 114 | 95 | 1565 | 1596 | 1579 | 1598 |
| $T_R$ | 147 | 138 | 133 | 92 | 1546 | 1555 | 1560 | 1601 |
| $T_O$ | 213 | 96 | 274 | 148 | 1431 | 1548 | 1370 | 1496 |

### 5.2. RQ2: Decoding Neuronal Dynamics Related to Impairment and Task Classification

From the perspective of cognitive impairment, we alter the response variable to address a 10-class cognitive task prediction problem, allowing us to expand the research question of “Does the depression of neuronal firing due to hypoxia lead to events of criticality with unique spectral arrangements?”, based on the tasks shown in Figure 5.

The 10-class model provides summary metrics that yielded an MCC of 0.54 for the *SS algorithmic strategy*, which was the worst-performing algorithm. The SI algorithm does slightly better with an MCC of .56, providing an increase of 3.57% in performance. The CSI approach provides an MCC of .60, which provides a performance increase of 7.14% and 11.11% compared to the SI and SS, respectively. The best-performing algorithm demonstrates superior discrimination with an MCC of 0.727, which is a 34.62% increase in performance compared to the worst algorithm SS and a 29.82% increase in performance to the traditional SI approach.

We provide Table 6 to further break down each algorithm’s performances using the TP, NP, FP, and TN for each of the ten classes. Similar to the 5-class model, the SIS algorithmic strategy is the best discriminator across all classes, except for the 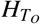 and 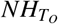. Thus, the discrimination performance using SIS seems similar to the 5-class model and has limitations with *T*_*O*_ as a whole. CSI remains the top-performing approach for distinguishing the specific class *T*_*O*_, regardless of whether it is a 5-class or 10-class model. Although the CSI algorithm had the second-best MCC when compared to the other approaches, the majority of the second-best TP classified featured are located within the SI algorithm strategy 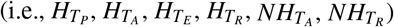. Regarding true positives and false negatives, SI, previously one of the weakest performers, has shown notable improvement and is now on the verge of becoming the second-best performer. SI’s performance hurdles and limitations are due to the performance in the false positive and true negative rates.

**Table 6:** This table compares all induced tasks and hypoxia conditions across the four algorithmic strategies, presenting the number of True Positives, False Negatives, False Positives, and True Negatives. The highlighted cells provide a visual comparison of performance: dark green indicates the best performance, light green denotes the second-best performer, orange marks the second-to-worst performer, and red highlights the worst performer. This coloring scheme aids in identifying which algorithmic strategy is most favorable for specific tasks.

|  | True Positive |  |  |  | False Negative |  |  |  |
| --- | --- | --- | --- | --- | --- | --- | --- | --- |
|  | SI | CSI | SS | SIS | SI | CSI | SS | SIS |
| $H_{T_P}$ | 130 | 126 | 120 | 145 | 82 | 86 | 92 | 67 |
| $H_{T_A}$ | 123 | 119 | 121 | 159 | 89 | 93 | 91 | 53 |
| $H_{T_E}$ | 120 | 105 | 108 | 163 | 92 | 107 | 104 | 49 |
| $H_{T_R}$ | 126 | 126 | 116 | 148 | 86 | 86 | 96 | 64 |
| $H_{T_O}$ | 162 | 215 | 152 | 184 | 68 | 15 | 78 | 46 |
| $NH_{T_P}$ | 112 | 110 | 113 | 148 | 87 | 89 | 86 | 51 |
| $NH_{T_A}$ | 124 | 111 | 111 | 151 | 75 | 88 | 88 | 48 |
| $NH_{T_E}$ | 113 | 107 | 123 | 146 | 86 | 92 | 76 | 53 |
| $NH_{T_R}$ | 119 | 119 | 111 | 152 | 80 | 80 | 88 | 47 |
| $NH_{T_O}$ | 148 | 223 | 164 | 190 | 82 | 7 | 66 | 40 |

Table 6: This table compares all induced tasks and hypoxia conditions across the four algorithmic strategies, presenting the number of True Positives, False Negatives, False Positives, and True Negatives.
|  | False Positive |  |  |  | True Negative |  |  |  |
| --- | --- | --- | --- | --- | --- | --- | --- | --- |
|  | SI | CSI | SS | SIS | SI | CSI | SS | SIS |
| $H_{T_P}$ | 82 | 94 | 89 | 33 | 1810 | 1798 | 1803 | 1859 |
| $H_{T_A}$ | 93 | 93 | 88 | 59 | 1799 | 1799 | 1804 | 1833 |
| $H_{T_E}$ | 66 | 71 | 53 | 52 | 1826 | 1821 | 1839 | 1840 |
| $H_{T_R}$ | 86 | 79 | 84 | 60 | 1806 | 1813 | 1808 | 1832 |
| $H_{T_O}$ | 97 | 37 | 120 | 60 | 1777 | 1837 | 1754 | 1814 |
| $NH_{T_P}$ | 92 | 78 | 66 | 61 | 1813 | 1827 | 1839 | 1844 |
| $NH_{T_A}$ | 74 | 76 | 87 | 65 | 1831 | 1829 | 1818 | 1840 |
| $NH_{T_E}$ | 66 | 61 | 74 | 30 | 1839 | 1844 | 1831 | 1875 |
| $NH_{T_R}$ | 85 | 87 | 79 | 42 | 1820 | 1818 | 1826 | 1863 |
| $NH_{T_O}$ | 86 | 67 | 125 | 56 | 1788 | 1807 | 1749 | 1818 |

## 6. Discussion

In earlier sections, we theorized, based on Newell’s timescale, that cognitive processes unfold across multiple temporal levels, and that analyzing EEG dynamics at granular, millisecond-scale intervals provides important information about the behavior of neuronal oscillatory groups. These dynamics can be captured through summated intensity, proportional shifts, stochastic fluctuations, or entropic measures that assess the stability of interactions among oscillatory components. Each of the four algorithms we introduced—Spectral Intensity (SI), Spectral Stability (SS), Competing Spectral Intensity (CSI), and Spectral Intensity Stability (SIS)—captures a distinct yet complementary aspect of these spectral dynamics. While previous work has largely focused on SI [24], our study introduces three new feature families—SS, CSI, and SIS—that aim to characterize how oscillatory activity changes over time during task-based cognitive processing. These new approaches go beyond traditional power analysis by quantifying not only the strength of specific frequency bands but also how their relationships shift, compete, and reorganize in response to task demands.

The entropic methods introduced in SS and SIS quantify the stability of spectral hierarchies by tracking how the relative ranking of frequency bands evolves over time. This provides a new perspective on how neural systems coordinate activity across multiple frequencies to support cognitive functions. ***We propose that these findings support a new conceptual model for cognitive state detection: Granular Analysis Informing Neural Stability (GAINS)***. GAINS highlights the idea that tracking fine-scale changes in spectral structure, rather than relying on coarse averages of power, offers more meaningful insight into the organization and dynamics of brain activity. We do not claim that our specific implementations, such as TCBE or the dynamic engagement index, represent the final or ideal solution. Instead, we view them as first steps in operationalizing the GAINS principle. Our broader goal is to motivate the field toward exploring how fine-grained spectral dynamics can improve EEG-based cognitive state classification.

More specifically, our proposed algorithms reflect the GAINS concept by examining how spectral intensities form evolving hierarchical patterns over time. These changes, illustrated in Figure 2, appear to encode task-relevant cognitive transitions. In simpler terms, the moment-to-moment self-organization of frequency bands seems to correspond with the functional demands of different tasks. To explore this idea, we presented four algorithmic strategies, each offering a unique view into EEG spectral activity. Together, these algorithms represent a progressive improvement in capturing informative features, incorporating granularity, temporal dynamics, and spectral relationships to enhance our ability to interpret brain activity.

The *task classification* and *cognitive impairment task classification* models demonstrated comparable predictive power between the traditional SI algorithm and the SS algorithm. It is important to note that these algorithms rely on fundamentally different bases of information. Spectral Stability measures hierarchical arrangements of frequency bands, quantifying how often specific arrangements occur and their predictability at the circuit level. This measure, catenated to the resolution of operations, serves as an entropy metric capturing the stability of firing pattern arrangements without considering the intensity of spectral bands. This concept of discriminative information is visually depicted in Figure 2, where the stability dynamics are highlighted in the outer figures. Consequently, SS provides granular contextual information about neuronal circuit dynamics on a temporal timescale of approximately 3.9 milliseconds (using a 256Hz sampling rate), illustrating how fundamental building blocks collaborate to form cognitive processes. The entropy measured by SS appears to be task-specific, resulting in a unique criticality arrangement for each cognitive task.

In contrast, Spectral Intensity focuses on summed intensities over time, reflecting operational and unit task resolutions derived from independent frequency bands without accounting for their interactions. Spectral intensity can be calculated through two methods: the standard approach, which sums the power over specific frequency ranges obtained from a Fourier Transform (illustrated in the central figure of Figure 2), and a secondary approach, which sums the intensity of the bandpass filter, 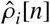, over time. The latter method captures circuit-level intensities but loses granularity through temporal summation, resulting in information aggregated at the operations or unit task level. The strikingly similar performance of SI and SS for both the 5-class and 10-class problems highlights the critical importance of both spectral intensity and its dynamics (hierarchical arrangements) over time.

Building on this, we introduce the Competing Spectral Intensities (CSI) algorithm, which integrates spectral intensity and dynamic information from spectral bands but differs from SS in two key ways: 1) employing a less granular timescale and 2) using a coarse measure of spectral dynamics. CSI is conceptually analogous to a coarse combination of SI and SS, summarized at the operations and unit task level using classical statistics (mean and variance). By merging spectral intensity and frequency band dynamics, CSI achieves enhanced predictive power compared to algorithms that rely solely on one form of information. For the 5-class problem, CSI achieves an MCC of 0.63, reflecting a 21.1% improvement over SI and a 16.6% improvement over SS, as shown in Figure 9. For the 10-class problem, CSI shows an 8.9% improvement over SI and a 12.9% improvement over SS, as seen in Figure 10. In Table 5, CSI emerges as the second-best algorithm for most tasks (3 out of 5) when comparing True Positives (TP) and False Negatives (FN) across all algorithms, occasionally tying with SI. However, for the 10-class problem (Table 6), SI outperforms CSI in TP rates for 6 out of 10 tasks.

**Figure 10:**
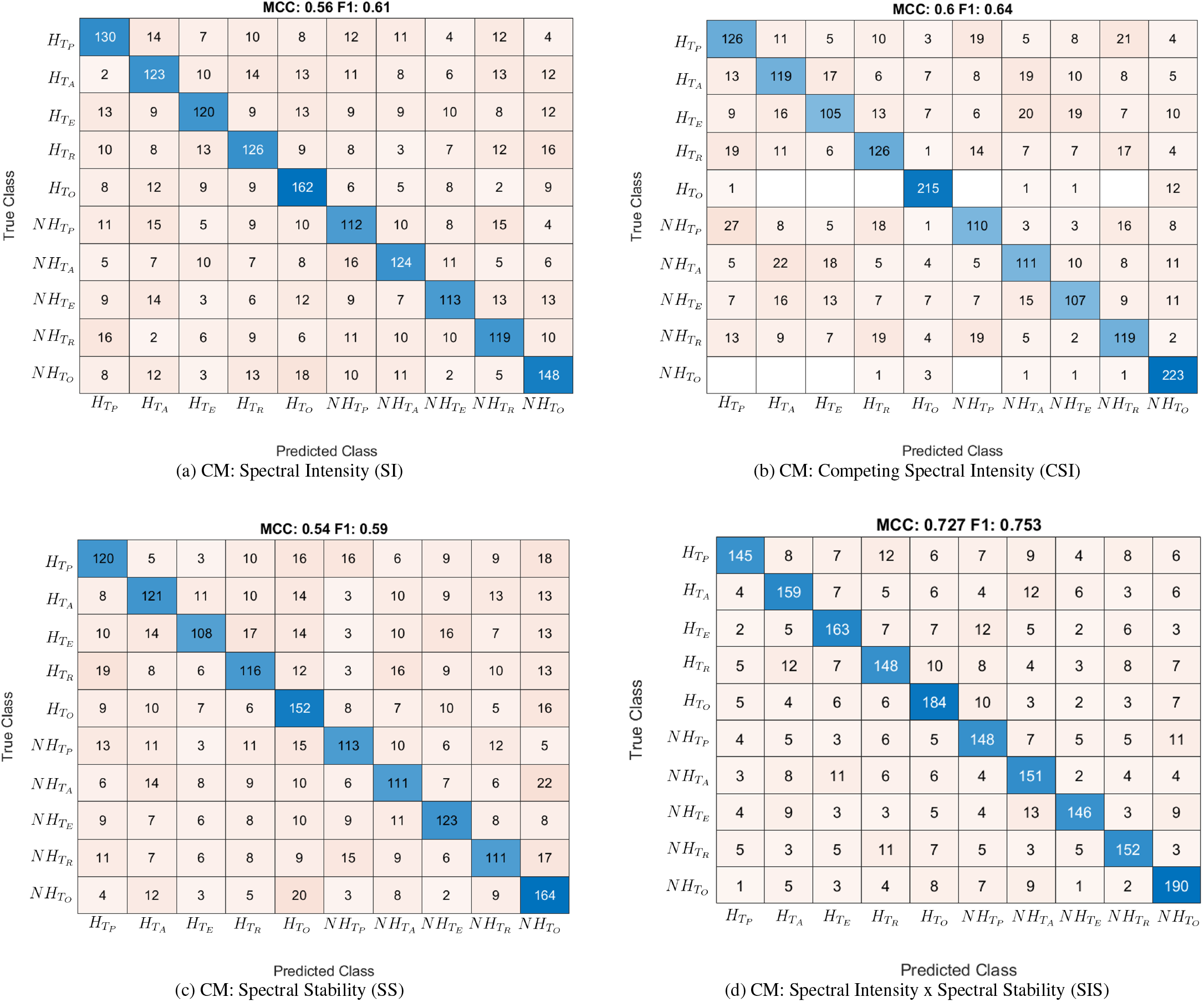
The four confusion matrices (CM) compare the predictive performance of the following methods: (1) the classical intensity analysis, which provides intensity information without incorporating temporal dynamics; (2) the proposed dynamic engagement indices, representing the stochastic nature of spectral intensity dynamics over time; (3) the proposed TCBE, which captures the complexity of spectral temporal dynamics while excluding intensity information; and (4) the interaction of TCBE with spectral intensity, representing refined temporal spectral intensity dynamics.

To address CSI’s limitations in temporal granularity and dynamic specificity, we developed the Spectral Intensity Stability (SIS) algorithm. SIS introduces a novel approach by combining instantaneous spectral intensity with per-sample entropy-based measures of spectral hierarchy, allowing it to track how the organization of frequency bands evolves over time. Unlike CSI, which summarizes intensity dynamics over broader time windows, SIS operates at the resolution of individual EEG samples (3.91 ms at 256 Hz), making it highly sensitive to fast, transient shifts in brain state. This sample-level precision, paired with a physiologically grounded filter bank, enables SIS to capture dynamic reorganizations in spectral structure that are often missed by traditional or windowed methods. SIS achieved the highest performance in both cohort-agnostic task prediction (30.7% improvement over SI, 25.9% over SS, and 7.9% over CSI) and cognitive impairment classification (29.8% over SI, 34.6% over SS, and 21.1% over CSI). These results suggest that SIS provides uniquely informative features for decoding cognitive state. Notably, its strong performance in distinguishing hypoxic from normoxic task conditions indicates a potential sensitivity to spectral reorganizations induced by oxygen deprivation. This opens a promising avenue for using SIS in real-time cognitive monitoring in high-risk environments such as aviation, deep-sea diving, or space missions. While this study did not directly examine the neurophysiological mechanisms underlying hypoxia, the SIS framework offers a practical tool for detecting subtle, task-relevant disruptions in neural organization.

Through this work, we identify three key factors for decoding cognitive state from EEG-derived global field potentials: (A) fine-grained temporal resolution (on the order of tens of milliseconds), (B) accurate estimation of spectral intensity across physiologically meaningful frequency bands, and (C) characterization of how these spectral components interact through evolving hierarchical arrangements. We consider these to be foundational dimensions, but additional features, such as transient waveform patterns and spatial source localization, may further enhance interpretability and performance. This study contributes to the broader discussion of EEG feature selection by proposing metrics designed to capture the dynamic organization of neural activity. By examining interactions across frequency bands and tracking the temporal reorganization of oscillatory patterns in response to task demands, these features offer a more precise and responsive framework for cognitive state classification.

## 7. Conclusion

This study establishes the concept of Granular Analysis Informing Neural Stability (GAINS), demonstrating that granular temporal interactions and hierarchical spectral intensity dynamics provide critical insights into neuronal firing patterns and their relationship to cognitive state and impairment. The Spectral Intensity Stability (SIS) algorithm outperformed others, showcasing the importance of combining spectral intensity and dynamic granular hierarchical arrangements for accurately decoding cognitive fingerprints and impairment, such as those induced by hypoxia. These three key factors—granular timescale resolution, spectral intensity, and hierarchical spectral dynamics—were identified as foundational for characterizing the neuronal criticality from these processes. While the approach is promising, further refinement and incorporation of transient behaviors and spatial dynamics are necessary for a more comprehensive understanding of neural self-organization and firing.

## Declarations

### Ethics approval

Approval for this study was granted by the Institutional Review Board of NASA Langley Research Center.

### Consent to participate

Informed consent was obtained from all individual participants included in the study.

### Consent for publication

Not applicable

### Availability of data and materials

The data that support the findings of this study are available from the corresponding author upon reasonable request.

### Code availability

The code that supports the findings of this study is available from the corresponding author upon request.

